# Windows to the goal: Pupillary working memory signatures prospectively adapt to task demands

**DOI:** 10.64898/2025.12.06.692784

**Authors:** Yueying Dong, Yun-chen Hung, Connie Xie, Anastasia Kiyonaga

**Author notes:** Senior author (Anastasia Kiyonaga).

## Abstract

The pupillary light response was once considered a brainstem reflex, but newer findings indicate that pupil dilation can also reflect content held ‘in mind’ with working memory (WM). This suggests that WM may recruit even the earliest sensorimotor apparatus for maintenance. Here, we tested two boundaries of this pupillary WM response: whether it generalizes beyond low-level stimuli and whether it adapts to changing behavioral goals. Namely, we tested whether the pupils reflect remembered brightness for real-world scene images, and whether the effect varies when different features dimensions are emphasized for the memory test (i.e., visual detail vs. semantic category). We found a feature-specific pupillary WM effect for remembering natural scenes, but only when the task encouraged a visual maintenance strategy. Rather than a retrospective echo of sensory-evoked stimulus features, the pupillary WM response prospectively adapts to how the memory content will be used.

## INTRODUCTION

In the opening minutes of the 1982 cinema classic *Blade Runner*, a Voight-Kampff machine zeroes in on the pupil of a test subject as he’s asked a series of evocative questions. In the audience, we understand that the pupillary response is meant to diagnose something deeper. Indeed, the plot device serves as a key determinant of whether the subject is a real human or a potentially dangerous replicant. The pupils have long been appreciated as windows into underlying states, and recent research has gone further to show that pupil size modulations might even reflect specific visual features of information that is held in mind^1–3^. Such pupillary signatures of working memory (WM) content stir the possibility that internal goal representations are expressed at the earliest stages of sensorimotor processing, and that the pupils are under more fine-grained cognitive influence than previously realized.

The pupillary light response (PLR) adapts vision to changing lighting conditions, constricting (or dilating) the pupils when more (or less) intense luminance hits the retina. This involuntary response was first considered a brainstem reflex, regulating light influx to the eye via photosensitive cells that project directly to subcortical targets^4–6^. Later, factors like arousal and cognitive load were found to evoke non-specific pupil dilation, likely via the locus coeruleus norepinephrine system^7–9^. Now, covert attention and mental imagery have been shown to conjure a stimulus-specific PLR-like effect under constant luminance^10–14^. For instance, the pupils become relatively dilated when simply imagining a darker stimulus (as opposed to a brighter one)^10,14^, suggesting that pupil signals also integrate with the more complex image-forming visual system. Therefore, while it had previously been known that pupil control might happen through well-characterized illumination-dependent pathways or state-based arousal pathways, feature-specific cognitive pupil modulations indicate that pupil control can happen through another pathway—one that is just now coming to light.

Pupil size can track the remembered brightness of stimuli maintained in WM, but the boundaries of this effect are undefined^1,2,15^. For instance, if both a dark and bright sample stimulus are initially encoded, and then the darker stimulus is retroactively cued as task relevant, the pupils will become relatively more dilated during the WM delay (vs. less dilated if the brighter stimulus is cued)^1^. This finding suggests the exciting possibility that the pupils provide a feature-specific index of the content held in mind. However, the functional importance of this pupillary WM response is unknown. For one, the effect has been shown for low-level visual stimuli – like simple black and white discs or oriented gratings of varying luminance^1,2,16^ – but it is unclear whether it generalizes to complex naturalistic images that vary on many more feature dimensions. Further, it is unclear whether the effect signals a retrospective response to item brightness, or a more functionally sensitive read-out of internal goal representations that guide upcoming behavior.

The observed pupillary WM effect is consistent with a sensory recruitment perspective, where WM shares circuits with sensorimotor processing^17–19^. In this framework, the pupils may reflect top-down activation of luminance codes in the visual cortex that propagate to pupil control pathways^5,20^. Yet, it remains contested what information is represented in sensory cortical activations during WM, and which underlying WM representations might link to pupil control^21,22^. One possibility is that the pupil signal recapitulates the sensory-evoked code, in which case the pupil response would closely mirror the physical features of encoded stimuli. An alternative is that pupil engagement supports prospective and action-oriented WM objectives, in which case the pupillary WM response should depend on anticipated behavioral demands^16,21^. Likewise, the strength and information content of neural WM activations can shift to reflect behaviorally relevant stimulus dimensions^23–29^. And the relative engagement of fronto-parietal vs. posterior sensory regions during WM has been shown to depend on whether categorical vs. precise visual stimulus features are more task-relevant^30^. If the pupils reflect a prospectively-oriented signal, they should exhibit similar flexibility to task context and behavioral goals.

In a prior study, we found that the magnitude of the pupillary WM response was greater for orientation stimuli that were most strongly prioritized^16^, hinting that pupillary WM signatures may adapt to behavioral context. Moreover, the effect was amplified in participants who reported having stronger visual imagery. Indeed, cognitive pupil modulations tend to vary with individual tendencies for visual or verbal thinking. Subjects who report weaker visual imagery show less pupillary response to imagined or remembered brightness^31^, and also report using more semantic-based memory strategies (rather than visual)^32,33^. Therefore, we reasoned that the pupillary WM response may depend on the representational strategies engaged during maintenance (i.e., visual vs. semantic). While previous work shows that cognitive pupil modulations correlate with trait imagery tendencies across individuals, here we aim to experimentally manipulate which task strategies are used within individuals. Specifically, we hypothesize that the pupils may only reflect remembered brightness when WM content is represented in a visual format. Here, we set out to test that possibility.

We asked three main questions: Does the feature-specific WM pupillary response emerge only for low-level stimuli, or do the pupils convey feature information about WM for complex natural scenes? Does the signal agnostically reflect the sensory features of WM content, or is it adaptive to behavioral goals? And how does the signal interact with an individual’s tendency for visual thinking, under different task goal conditions? To answer these questions, we had participants encode both daytime and nighttime scene images for a delayed recognition task. We manipulated which item would be tested as well as which stimulus dimension (visual vs. semantic) would be emphasized for the test. If the pupillary WM response is goal-oriented and sensitive to different maintenance strategies, we would expect it to be relatively enhanced under more visual task goals, and relatively diminished under more semantic task goals. Together this work will illuminate the boundaries of pupillary WM and the pathways by which the eyes reflect our inner thoughts.

## RESULTS

### Visual vs. semantic cues prompt distinct performance profiles

Participants (*n* = 44) completed a delayed recognition task for real-world scene images (**Figure 1A**). On every trial, two WM samples were shown: one daytime scene (i.e., brighter) and one nighttime scene (i.e., darker; **Figure 1B**). After encoding, a directional cue (‘retrocue’) indicated which of the two samples would be tested at the end of the trial. The color of the retrocue also indicated which dimension of that sample would be tested – visual detail or higher-level object category. We term these trial conditions *visual* and *semantic*, respectively, for shorthand. At test, participants selected the cued item from a probe array (**Figure 1C**). In the *visual* condition, they had to select the probe that matched the exact cued exemplar. In the semantic condition, they had to select the probe that matched the cued item category (see ***Stimulus image processing*** for all category labels). The probe arrays contained condition-specific lures designed to encourage prioritizing the cued stimulus dimension. In theory, the visual condition should require a more perceptually detailed representation, to discriminate the sample from semantically-similar lures. The semantic condition, on the other hand, might encourage a verbal label to discriminate the cued category from semantically-distinct but visually-similar lures. In short, given the same encoded content, we aimed to manipulate how that content was maintained.

**Figure 1.**
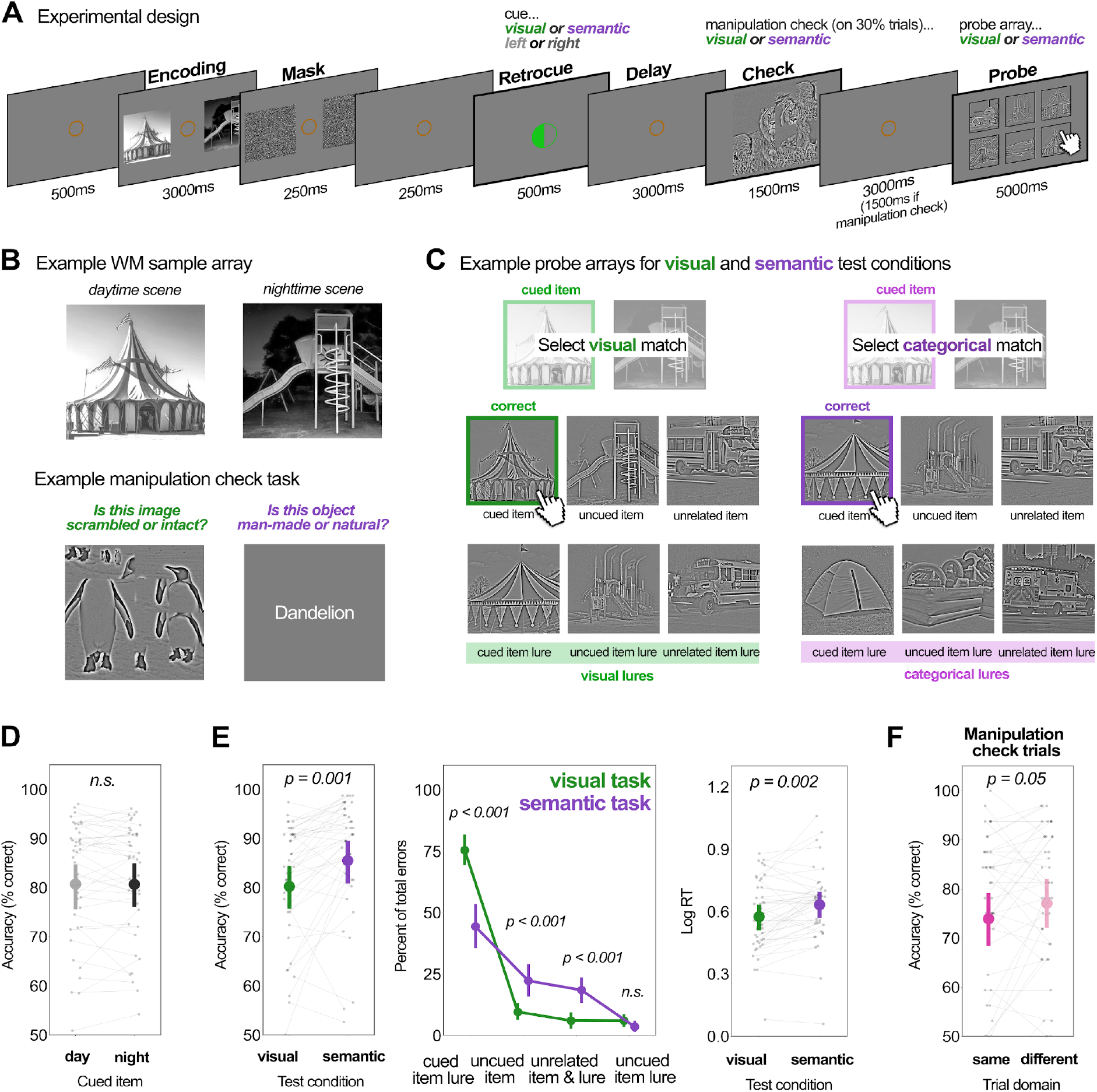
Experimental design and behavioral performance. (**A**) Task schematic for an example trial. The WM sample array on each trial consisted of a day- and nighttime scene. After encoding, a color cue indicated the to-be-tested WM sample stimulus, as well as the relevant stimulus dimension for the probe (visual detail or semantic category). During the delay, a manipulation check sometimes occurred, during which participants made either a visual or semantic judgement. After the delay, participants were probed to select the correct image from an array of six options. (**B**) Example WM samples (top) and manipulation check stimuli (bottom). (**C**) Example probe arrays for visual and semantic trials. Each array contained matches for both the cued and uncued items, lure images for the cued and uncued items, and 2 items from an unrelated category (unrelated lures). On *visual* trials, the cued item lure was another item from the same semantic category (e.g., if a circus tent was cued, a different circus tent would also appear in the probe array). On *semantic* trials, the cued item lure was a visually similar item from a different semantic category (e.g., if a circus tent was cued, a camping tent would also appear in the probe array). Probe stimuli were high-pass filtered to remove luminance information. (**D-F**) Behavioral performance. Small data points reflect individual subject means; bolded points reflect the group mean; error bars reflect the 95% CI. **(D)** WM choice accuracy (% correct) for daytime (grey) and nighttime (black) images. (**E**) WM choice accuracy (left) and RT (right) for *visual* (green) and *semantic* (purple) task conditions (on trials with no manipulation check). Log RT reflects the log transformed duration between probe onset and mouse click. The middle panel shows the percentage breakdown across the possible types of erroneous response. (**F**) WM choice accuracy on manipulation check trials, as a function of whether the intervening task taxed the same (darker) or different (lighter) stimulus dimension as the WM task. The significance value marks a main effect of the manipulation check domain congruency (i.e., same vs. different from the WM task).

We also incorporated an occasional manipulation check – in the form of an intervening visual or semantic judgment task – to verify whether participants prioritized the expected stimulus dimension (**Figure 1B)**. Note that the task-relevant item and stimulus dimension were both cued after encoding, so that any pupillary differences between conditions could be attributed to WM maintenance of the selected item, rather than any sensory or attentional differences from encoding (see ***Methods*** for more detail).

Performance was good overall (mean accuracy: 82.87%), and paired-sample *t*-tests confirmed that memory did not differ between daytime versus nighttime scenes (**Figure 1D**; accuracy: *t(43)* = 0.10, *p* = 0.91, Cohen’s *d* = 0.00,; RT: *t(43)* = 0.31, *p* = 0.76, Cohen’s *d* = 0.01). Therefore, on trials uninterrupted by a manipulation check, it is unlikely there was a difficulty difference between remembering the darker or brighter stimuli.

However, participants did display different performance profiles for *visual* vs. *semantic* task conditions. WM probe choices were more accurate when the semantic category was probed, as compared to visual detail (**Figure 1E**, left; *t*(43) = -3.63, *p* = 0.001, Cohen’s *d* = -0.36). Conversely, responses were faster when visual detail was probed, as compared to semantic category (**Figure 1E**, right; *t*(43) = 3.36, *p* = 0.002, Cohen’s *d* = -0.33). This hints that, given the same encoded content, the retrocues may have prompted different task strategies that alter either the quality or accessibility of the relevant content. Indeed, follow-up analyses also revealed different error profiles between the two test conditions (**Figure 1E**, middle). On *visual* trials, errors were made overwhelmingly toward a lure image from the same category as the target item. On *semantic* trials, however, participants were relatively less likely to make a comparable error toward a visual lure (*visual* vs. *semantic*: *t*(43) = 6.71, *p* < 0.001, Cohen’s *d* = 1.29), and more likely to instead choose the uncued sample item (*t*(43) = -3.83, *p* <0.001, Cohen’s *d* = -0.79) or an unrelated item (*t*(43) = -4.46, *p* <0.001, Cohen’s *d* = -0.91). Thus, the *visual* vs. *semantic* cues may have prompted different maintenance strategies that altered the specificity of the WM representation.

The manipulation check (30% of trials) further corroborated that participants were using different maintenance strategies in the two conditions. This check comprised an intervening task that required either a visual or semantic judgement during the WM delay. The visual judgement was to decide whether an image was scrambled or intact, and the semantic judgement was to decide whether a text string described a natural or manmade object (**Figure 1B**, bottom panel). If participants were using the retrocues as expected, WM performance should be worse when the manipulation check taxed the same stimulus dimension as the WM test^30^. We found this to be the case (**Figure 1F)**. A repeated measures ANOVA with factors of WM cue condition (*visual* vs. *semantic*) and manipulation check domain (same vs. different) found both a main effect of WM condition on accuracy (mirroring the *t*-test above; *F*(1,43) = 7.39; *p* =0.01 ; *η*_*p*_^*2*^ = 0.15), as well as a main effect of manipulation check domain congruency (*F*(1,43) = 4.06; *p* = 0.05; *η*_*p*_^*2*^ = 0.09). WM accuracy was relatively worse when the manipulation check was in the same domain as the WM cue condition. That is, WM was worse on *visual* trials under a visual manipulation check (vs. semantic), and worse on *semantic* trials under a semantic manipulation check (vs. visual). There was no interaction between WM cue condition and manipulation check domain (*F*(1,43) = 0.76; *p* = 0.39; *η*_*p*_^*2*^ = 0.02), indicating that the *visual* and *semantic* WM conditions were equally susceptible to same-domain disruption.

To summarize the behavior, participants were more accurate on *semantic* trials (vs. *visual*) but faster on *visual* trials (vs. *semantic*), hinting that the different behavioral goals may prompt distinct maintenance strategies. Likewise, the two conditions yielded different error profiles, and each condition was more susceptible to a competing task in the same domain. Therefore, participants appear to use the retrocues as expected, prospectively modulating their WM maintenance strategies in accordance with anticipated upcoming demands.

### Pupils express remembered brightness for complex real-world scenes

Prior work suggests that the pupillary response may provide a sensitive readout of the information content held in mind. Namely, the pupils relatively dilate when a darker (vs. brighter) item is maintained in WM, suggesting that WM content maintenance may extend to the earliest sensorimotor apparatus^1,2^. However, this ocular modulation has only been shown for low-level visual WM stimuli, like oriented Gabors. Our first pupillary analyses aim to gauge whether the feature-specific pupillary WM response replicates for multi-faceted real-world scene images.

Indeed, we found that pupils were larger during the WM delay when a nighttime stimulus was remembered (vs. daytime; **Figure 2a**). We used a cluster-based permutation *t-*test to compare the pupil time series after a daytime vs. nighttime scene was flagged by the retrocue. We found that the two pupil traces diverged later in the WM delay (2600ms to 3500ms after retrocue onset; *p = 0*.*023*). Note that probe images were high-pass filtered to remove their luminance, so that any pupil effects during the delay could be attributed to maintenance processes rather than probe expectations. This effect was also specific to trials where participants responded correctly (2600ms to 3300ms, *p = 0*.*033*), whereas there was no pupil size difference between brightness conditions on incorrect trials (no clusters detected; bottom row of **Figure 2a**). Therefore, the WM pupillary response that has previously been shown for low-level visual stimuli also occurs for memories of complex naturalistic scenes. The feature-specific pupillary WM response is further associated with successful WM maintenance, not a reflexive reaction to item cueing or driven by luminance conditions at probe.

**Figure 2.**
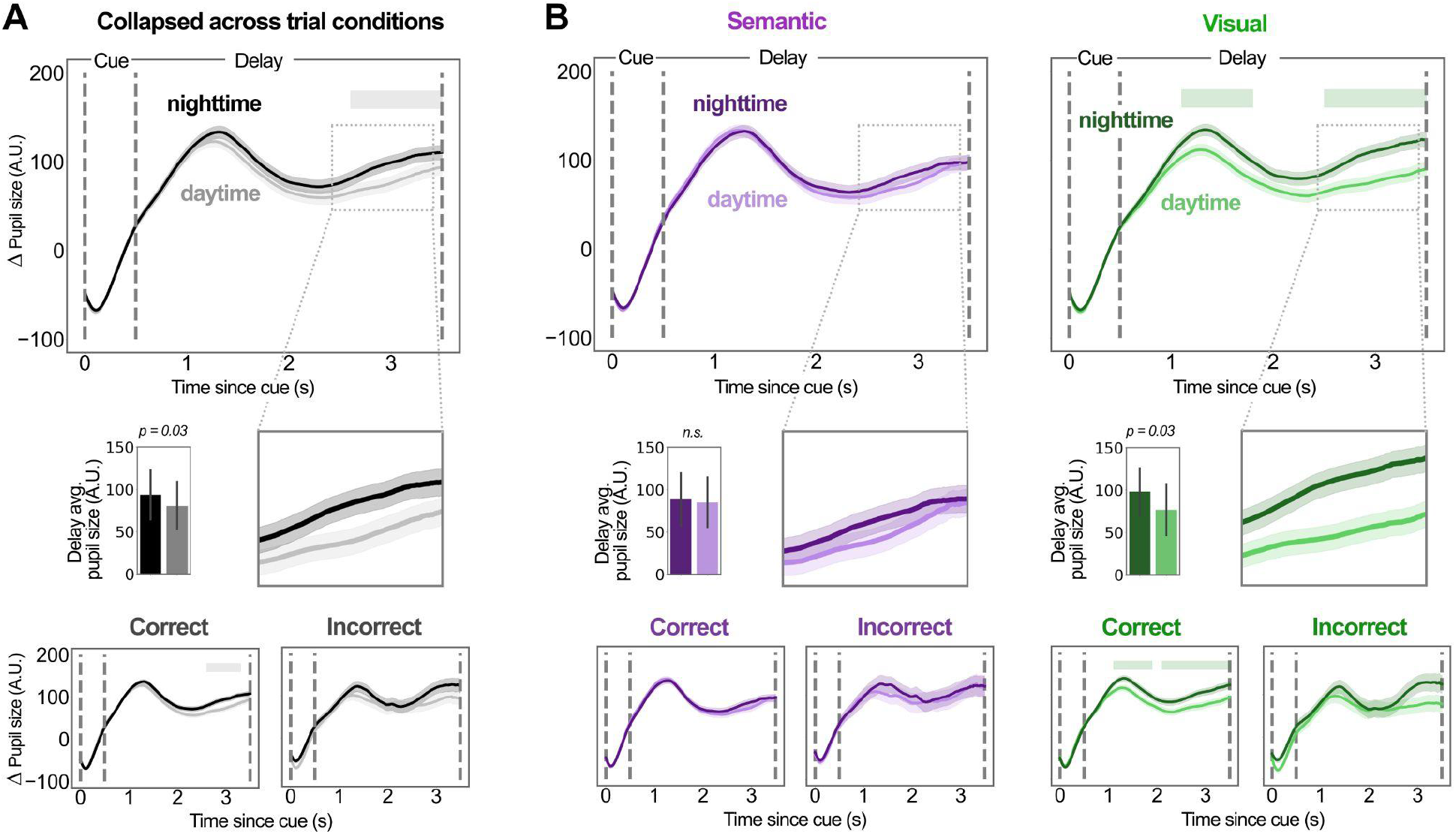
Pupil size timecourse across the working memory delay period. Upper panels show the grand average pupil size (baseline-normalized), in the period after the retrocue onset, for daytime (lighter shades) and nighttime (darker shades) scene images. The vertical dotted lines indicate event onsets, corresponding to retrocue, delay period, and manipulation check, respectively. The shaded horizontal bands at top of plot indicate periods where permutation testing yielded a significant difference between daytime and nighttime conditions. In the middle row, barplots depict the window-averaged pupil size over the entire delay epoch, for daytime and nighttime conditions. The bottom panels show the grand average of correctly (left) and incorrectly (right) responded trials. Across all panels, shaded error bands reflect ±1 SEM, and error bars reflect 95% CI. (**A**) Pupil timecourses collapsed across *visual* and *semantic* trial conditions. (**B**) Separate pupil timecourses for *semantic* (left, purple) and *visual* (right, green) trial conditions.

### Pupillary WM response occurs when visual detail is required

A primary goal of this study was to test whether the pupils either retrospectively track the sensory features of encoded content or prospectively signal how the content will be used. Specifically, we ask whether the pupillary WM response is flexible to the visual vs. semantic WM task demands. If the pupils reflect a coarse brightness tag that is agnostic to behavioral goals, then *visual* and *semantic* trial conditions should both show a comparable pupillary effect during the delay. If instead the pupillary WM response is specific to when a visual format of representation is engaged, then it should only be evident on *visual* trials (but not *semantic*).

Consistent with the latter hypothesis, we found that the pupillary WM response (nighttime - daytime scene) was only present when the WM test required memory of visual detail (vs. semantic category; **Figure 2b**). Permutation *t*-tests found no significant clusters of difference in pupil size between daytime and nighttime scenes for the *semantic* condition. However, for the visual condition, two clusters emerged: one shortly after cue offset and one later in the WM delay (time after retrocue onset: 1100ms to 1800ms, *p = 0*.*032*; 2500ms to 3500ms, *p = 0*.*016*). Paired-sample *t-*tests on the individual-level pupil size, averaged across the delay, corroborated the time series statistics (*semantic*: *t*(43) = -0.478, *p* = 0.635, Cohen’s *d* = -0.040; *visual*: *t*(43) = -2.203, *p* = 0.033, Cohen’s *d* = -0.210). To assess the evidence for the null hypothesis in the *semantic* condition, we also computed a Bayes factor on the individual-level pupil difference, which suggested moderate evidence for the null *(BF*_*01*_ = 5.49). Like the aggregated data (**in Figure 2a**), the condition-specific effect was only evident on trials when participants responded correctly (bottom row of **Figure 2b)**. Two significant clusters emerged only for correct *visual* trials (1100ms to 1900ms, *p = 0*.*035*; 2100ms to 3500ms, *p = 0*.*015*), but none for incorrect trials. For *semantic* trials, no significant clusters emerged regardless of correctness.

This result indicates that the pupils adaptively reflect WM maintenance strategies that vary trial-by-trial. Note that the probe conditions were retrocued after viewing the WM samples, so encoding demands were always matched between conditions, both in terms of the sensory-evoked brightness and the relevant stimulus dimension for the test (visual vs. semantic). Thus the observed pupillary effect emerges only during the WM maintenance process. A brightness-related pupillary WM response may only arise when people engage a representation that requires visual detail rather than a high-level category label.

### Strong imagers exhibit pupillary WM response to memories of semantic categories

Individuals who report more vivid visual imagery tend to have a larger cognitively modulated PLR-like effect, while those with weaker imagery have a small or absent effect^33^. This cognitive PLR has therefore been proposed as a physiological marker of the integrity of an individual’s visual mental representations^10,14,31^. Here, in an exploratory correlational analysis, we ask how such self-reported imagery tendencies interact with instructed task strategies.

We tested whether an individual’s pupillary WM response (difference in remembering nighttime - daytime scenes over the delay epoch of interest) correlated with their self-reported visual imagery strength (measured with VVIQ^34^). We found a positive correlation between the two for the *semantic* test condition (*r* = 0.335, *p* = 0.02), but not the *visual* test condition (*r* = -0.146, *p* = 0.342; **Figure 3b**). This experiment was not designed to test between-group effects, but we split the sample into three imagery bins (relatively weaker, moderate, or stronger imagers) to illustrate the nature of the correlation. We plot the pupil time series separately for each bin just for visualization purposes (**Figure 3c**). In the *semantic* condition, the pupillary WM response appears relatively larger for individuals with stronger visual imagery (compared to weaker or moderate imagery). In the *visual* condition, however, the pupillary WM response appears comparable for individuals across the imagery spectrum (at least the portion of the spectrum sampled here). Thus, we speculate that individuals with stronger imagery may engage visual strategies even when the task does not require it (i.e., *semantic* condition). Moreover, those who report relatively weaker imagery may be able to engage visual strategies as needed when instructed (i.e., *visual* condition). This variation across individuals highlights that reliance on visual or semantic WM strategies – and the accompanying engagement of pupillary pathways – may lie along a continuum that flexibly varies with intrinsic individual tendencies and behavioral demands imposed by the task.

**Figure 3.**
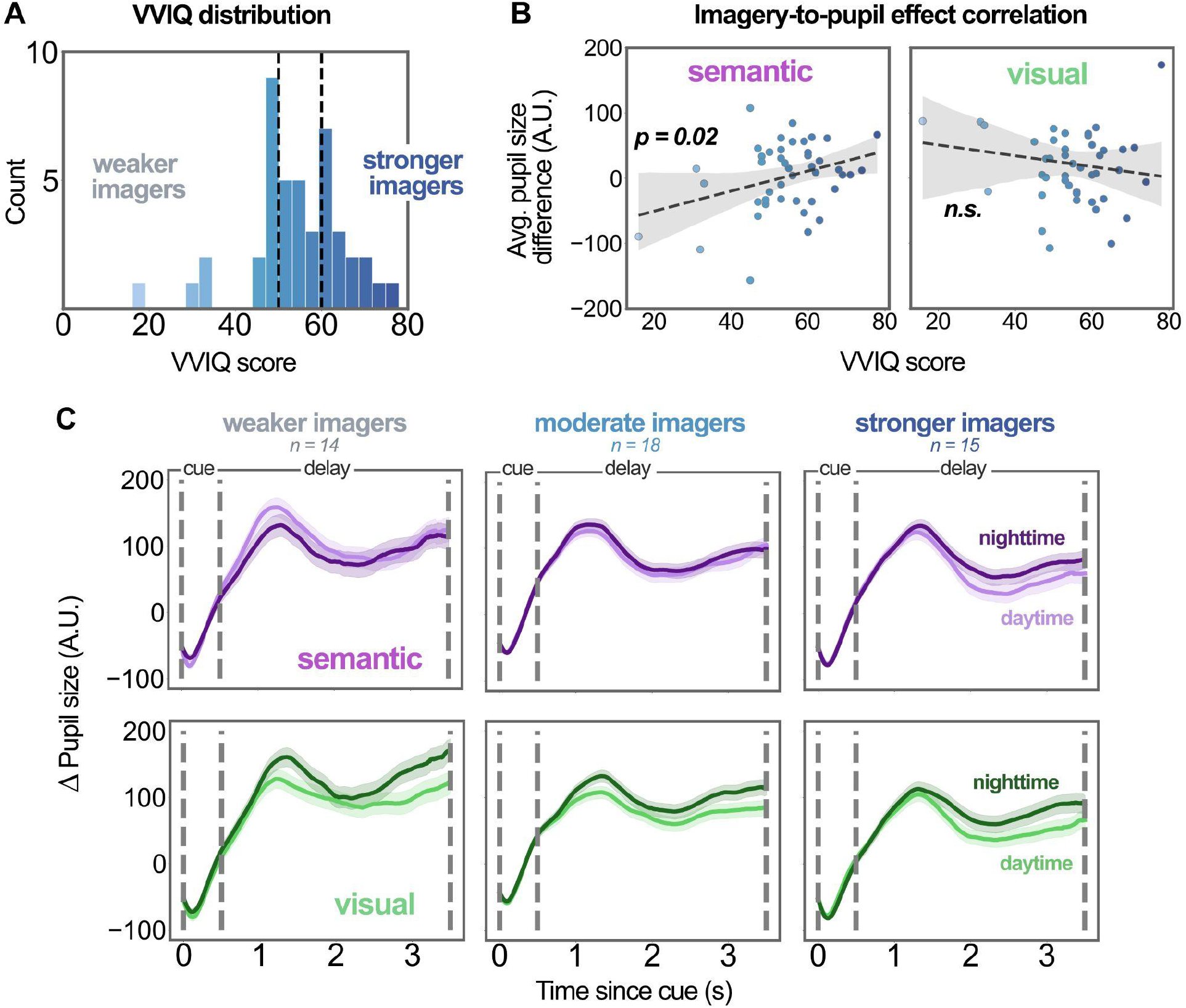
Individual differences in visual imagery and pupillary WM response. (**A**) Distribution of scores on self-report VVIQ questionnaire. Dashed lines mark the first and second tertiles (⅓) that partition participants into weaker, moderate, and stronger imagery groups for data visualizations in C. (**B**) Correlations between an individual’s VVIQ score and their magnitude of pupillary WM response (average pupil size for remembering nighttime - daytime scenes), for both *semantic* and *visual* task conditions. Dashed lines are linear fits to the data. (**C**) Grand average pupil timecourses, for remembering daytime and nighttime images, plot separately for each imagery group (weaker, moderate, and stronger imagers). Upper panels show the *semantic* condition (purple) and lower panels show the *visual* condition (green). Shaded error bands reflect ±1 SEM.

### Flexibility in pupillary working memory is distinct from eye movements and mental effort

Lastly, we address possible confounds that could explain the observed feature-specific pupillary WM response. Following from sensorimotor accounts of WM^17,18,35^, we theorize that the response accompanies recruitment of visual cortical representations during WM, which would be greater when more visual detail is required for the task. However, the basic brightness effect could theoretically emerge if darker stimuli were more cognitively taxing to remember. Likewise, the modulation by task condition could emerge if the *visual* vs. *semantic* conditions provoked different degrees of cognitive effort or attentional allocation^36–39^. Indeed, *semantic* trials were more accurate while *visual* trials were faster, which could occur if the trial types were differently demanding. Here, we examined two additional indices that should vary with attentional load – microsaccade frequency and index of pupillary activity (IPA) – to test whether they also varied with the relevant stimuli and WM task conditions.

Microsaccades are miniscule eye movements that are also associated with the allocation of attention in mnemonic visual space^40–43^. For instance, during a WM delay, microsaccades tend to veer toward the location where a relevant WM item was encoded, potentially indexing attentional selection within WM^44–48^. The frequency of such biased microsaccades is also greater toward more behaviorally relevant WM content^16^, further suggesting it indexes attentional deployment. If participants allocated more attention to darker stimuli or during the *visual* task condition, thereby explaining the relatively amplified pupillary response, we might also expect to see larger microsaccade biases in these conditions. However, we found no such differences. After the retrocue, we did find that gaze location was biased toward the visual hemifield of the task-relevant item, and microsaccade frequency increased in the same direction (**Figure 4A and B**). However, permutation *t-*tests on the time series found no significant differences in microsaccade frequency between either daytime vs. nighttime scene conditions, or *visual* vs. *semantic* cue conditions (no clusters detected). This suggests comparable attentional deployment across conditions.

**Figure 4.**
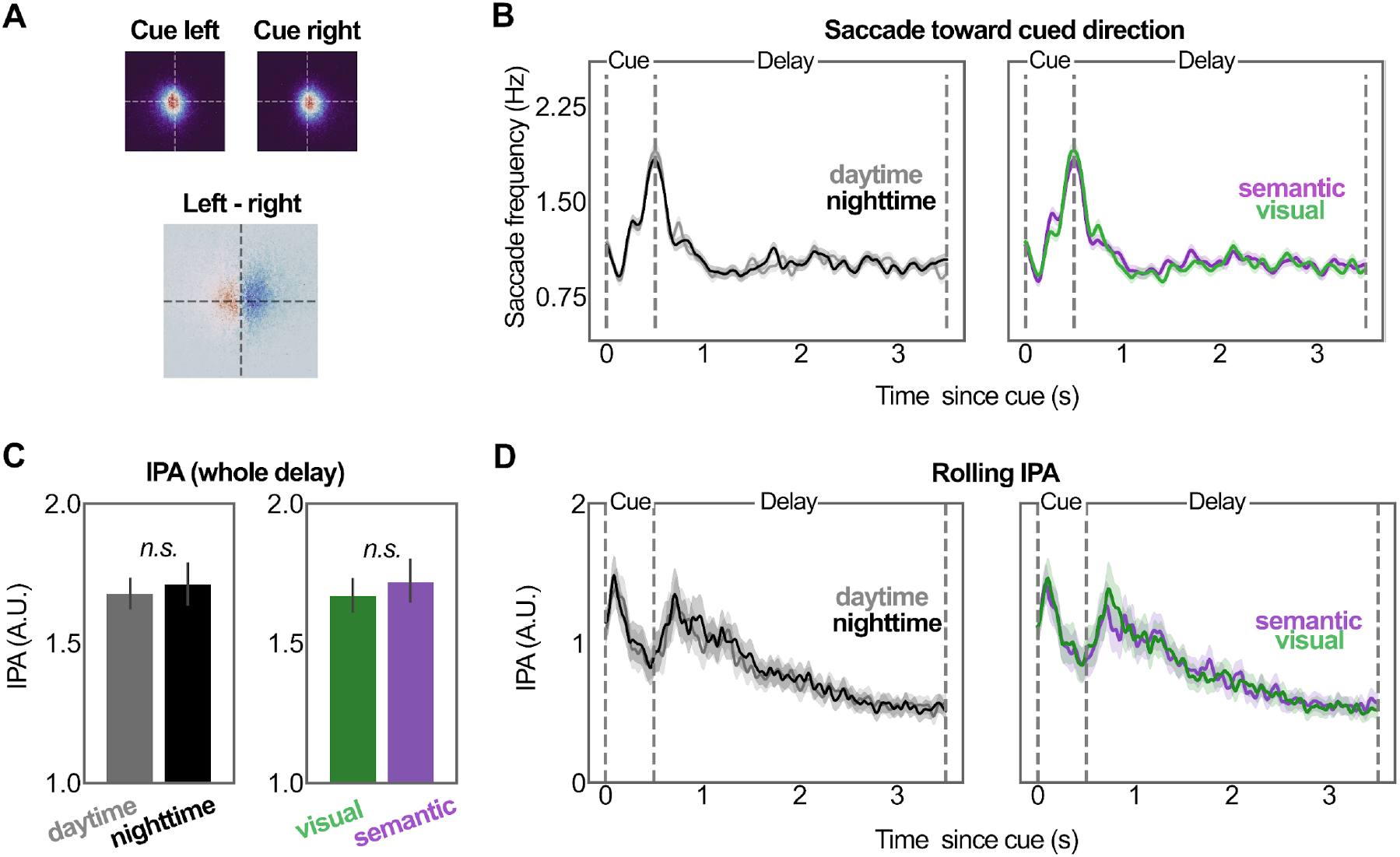
Saccade biases and index of pupillary activity (IPA). (**A**) Left: Gaze heatmaps for trials when the sample from either the left or right side of the encoding screen was cued, collapsed across time. Bottom panel shows the difference between left- and right-cued trials, and warmer color represents more gaze visitations at that location. (**B**) Frequency of saccades toward the cued direction (relative to uncued), across the trial. Plotted for day-vs. nighttime scenes (left) and *semantic vs. visual* trial conditions (right). (**C**) IPA scores (calculated from the whole delay period) for remembering daytime vs. nighttime scenes images (left) and for *visual* vs. *semantic* trial conditions (right). (**D**) Rolling IPA scores (100ms window) across the trial (after cue onset). Across all panels, shaded error bands reflect ± 1SEM and error bars reflect 95% CI.

The IPA measures oscillatory activity in the pupil timeseries. This index is thought to reflect cognitive effort and is unaffected by brightness-evoked changes in pupil size^36^. If either the darker WM stimuli or *visual* test condition was more effortful, thereby explaining the condition-specific pupillary WM response, we would expect a higher IPA score in those conditions. However, we found no such differences. A two-way ANOVA on the IPA, calculated over the whole 3-second delay, found no effect of cued item brightness (**Figure 4C**; *F*(1,43) = 1.914, *p* = 0.174, *η* _*p*_^*2*^ = 0.001), stimulus dimension conditions (*F*(1,43) = 0.431, *p* = 0.515, *η*_*p*_^*2*^ = 0.001), or an interaction between factors (*F*(1,43) = 0.033, *p* = 0.856, *η*_*p*_^*2*^ < 0.001). Time series analyses on IPA across the trial, using a 100ms rolling window, also confirmed no differences between either the brightness conditions or cued stimulus dimensions (no clusters detected; **Figure 4D**). There was an overall increase in IPA to the cue onset and offset, roughly around 150ms and 750ms respectively, but no evidence that the different task conditions evoked different degrees of cognitive effort (as measured by the IPA).

Taken together, the feature-specific pupillary WM response appears to index a distinct process from gaze biases during the WM delay, and to emerge independent of task difficulty.

## DISCUSSION

Here, we tested WM for complex daytime and nighttime scenes, and we manipulated whether precise visual detail or semantic category was behaviorally-relevant for the WM probe. We found that pupil size reflected remembered scene brightness during WM maintenance, especially when the task would require finer visual detail (vs. a category label). Therefore, the pupils convey a feature-specific WM content signal, but it flexibly varies with the underlying maintenance strategy: pupils preferentially reflect WM representations that are prioritized in a visual format. Rather than a retrospective echo of the encoded sensory stimulus features, the pupillary WM response prospectively adapts to how the memory content will be used.

### Pupillary working memory is cognitively flexible

These findings reveal fine cognitive nuance in the pupil. Recent work has shown that the brightness-related pupillary WM response is amplified for stimuli that are most strongly prioritized^16^, and here we show that it is additionally modulated by which stimulus dimensions are most task-relevant. The signal is therefore sensitive but selective, scaling with item relevance but seemingly unique to WM content held in a visual format. Given the same encoded content, cortical representations during WM can transform to reflect the prioritized feature dimension^21^, and here we see that such a transformation is also evident in the pupil signal.

We further show that the link between WM and pupillary control goes beyond simple low-level stimuli. There is some evidence that cardinal orientations have unique effects on cognitive pupillometry, raising concerns that the pupillary WM response may be a byproduct of specific stimulus effects^49^. Here, WM for complex real-world scene stimuli also evoked a brightness-related effect in the pupils, showing that the pupillary WM response generalizes to higher-dimensional stimuli. Note that the relevant stimuli and feature dimensions were cued after encoding here, and luminance information was stripped from the probe display images. Therefore, any pupil size differences between conditions should stem from how the information is maintained in WM. The effect also emerged only in correct trials, suggesting it precedes successful behavior. Therefore, in keeping with an expansive version of the sensory recruitment framework^19,21,35^, the visual representations that support WM-guided behavior may be selectively expressed as early as the sensory receptors.

### Ocular activity provides multiple selective processing indices

In this and other related work^16,43^, pupil size and eye movements show distinct patterns of flexibility to WM conditions. Here, the brightness-related pupillary WM response was sensitive to instructed WM strategy, whereas neither microsaccade biases nor a pupillary index of cognitive effort (IPA) showed such sensitivity. This suggests, for one, that the feature-specific pupillary WM response is not explained by attentional deployment or task difficulty^36,42,44^. Additionally, it indicates that the eyes can convey multiple unique indices of WM content and internal attention. For instance, brightness-related pupil size effects may index content information about what is maintained and in what representation format. Gaze and microsaccade frequency may convey spatial information about where internal attention is allocated and what locations are selected for processing^40–42,44–46^.

Within the feature-specific pupillary WM response, moreover, there may be indices of multiple distinct processes unfolding over time. Namely, in this and other related work^16^, at least two prominent pupil inflections emerge during the WM delay. There are periods of elevated dilation both shortly after the retrocue and again as the probe approaches, and both show a brightness-related effect. However, there are hints that these inflections may represent dissociable underlying function^50–52^. For instance, they show different patterns of sensitivity to attentional modulation and representational format^16^, and here the earlier peak and later rise show distinct relationships with individual imagery strength. In our data, the initial peak follows a sharp increase in biased microsaccade frequency, consistent with the idea that it reflects the content of attentional selection within WM^42,48,53^. Later in the delay, there was minimal microsaccade activity and therefore no evidence for a microsaccade marker of attentional selection. Instead, the late delay rise is consistent with preparatory activity for the probe, consistent with the neural ramping activity that is often observed leading up to a WM response^54–56^.

Beyond luminance, the pupils are known to respond to other visual features like numerosity, contrast, and proximity^57–59^. It is unknown whether such features can modulate the pupils endogenously during WM, but it seems likely, given the range of seemingly ‘perceptual’ effects that have been shown to emerge during WM^60–62^. The pupils therefore hold great promise to read out underlying representations at finer levels of granularity than previously realized.

### Representational states and traits for working memory

WM-related signals are detected in widely distributed regions across the brain^63,64^. But it is unclear which of these signals contribute unique WM functions, and which functions are reflected in ocular WM signatures. Some WM-related activations in visual cortex are thought to be explained by lingering sensory-evoked activation^65,66^, or a stable retrospective trace of encoded content^67^. A simple interpretation construes pupillary WM effects as an artifactual read-out that similarly recapitulates the sensory-evoked response. In that case, we would expect the pupil modulation during WM maintenance to closely follow the encoded stimulus features. However, many other WM-related neural signals appear prospective in nature, shifting in amplitude and content to meet anticipated task demands^21,24,68^. The current findings show that pupillary WM is modulated by expected demand, suggesting it aligns with such context-sensitive neural representations.

What does this suggest about how WM is carried out under different demands? WM may engage different underlying systems when different feature dimensions are prioritized. For instance, when specific featural detail is required for the task, participants may prioritize a high-resolution representation in visual cortex that also engages the pupils to reinstate the sensory stimulus details. This precise representation may sometimes be unsustainable and subject to interference from visually-similar lures, but highly-accessible for fast responding (matching our observed behavioral response profile in *visual* conditions). On the other hand, when a category label is more important than the visual appearance, it would be metabolically inefficient to reinstate the sensory stimulus details and pupillary circuitry during WM maintenance. A coarser categorical code might be engaged, via later semantic processing regions, which is less susceptible to visual interference but slower to retrieve at probe (matching our observed behavioral response profile in *semantic* conditions). However, both our *visual* and *semantic* tasks could potentially be completed with multiple different strategies, and several types of representation might support WM in either condition. For instance, the semantic condition might encourage verbal labelling or instead a coarser visual category representation. Our data are ambiguous as to whether the *visual* vs. *semantic* conditions invoke fundamentally distinct representational states vs. different degrees of visual cortex activation. But, either way, shifting between strategies likely lies along a continuum.

Indeed, individuals continuously vary in their reported tendency to engage visual or semantic representations in daily life^69^. Weaker imagery has been associated with reduced functional connectivity between visual areas and high-order control networks^70–72^. These connectivity differences may underlie the subjective experience of imagery as well as differences in pupil engagement^73,74^. Cognitive pupil effects are typically amplified in those who report strong visual imagery and dampened in those who report weak imagery^31^, but our data suggest these tendencies may also be flexible to task demands. The current sample was not powered to formally examine individual differences, so should be interpreted with caution, but the results suggest that instructed strategies may modulate intrinsic imagery tendencies in some cases. For instance, weaker imagers may typically default to a verbal labeling or a coarser pictorial representation that does not meet the threshold to engage brightness-sensitive pupil pathways^31,33,75^. However, when visual detail was cued as important here, the pupillary WM effect appeared comparable for weaker and stronger imagers. Therefore, those who report weak imagery may be able to engage visual representations and pupil control pathways^75^ when instructed. On the other hand, when a category label was prioritized, stronger imagers appeared to still show a brightness-related pupillary effect (compared to weaker and moderate imagers). Therefore, those who report stronger visual imagery may default to engage visual representations even when they are inessential to the task. Future work should seek to more extensively sample WM pupillary effects across the individual imagery spectrum while collecting more objective markers of potentially distinct cognitive strategies and associated representational states. Rather than a dichotomous predictor, pupillary WM may flexibly index the relative engagement of perceptually rich representations in visual cortex.

### Conclusions & implications

Here, we found that the pupils sensitively express the visual content that is held in mind. This work has both practical and theoretical implications for how we study and understand WM function. In practical terms, this highlights the value of pupillary and other ocular signals as refined indices of mental content. In theoretical terms, this illuminates the vastly distributed physiological networks that carry out WM. The findings begin to describe the boundaries of cognitively-modulated pupil pathways, and highlight the prospective nature of WM representations. Sensorimotor recruitment for WM may extend as early as the sensory receptors, where the pupils themselves are an instrument of flexible WM.

## RESOURCE AVAILABILITY

### Lead contact

- Requests for further information and resources should be directed to and will be fulfilled by the lead contact, Yueying Dong.

### Materials availability

- This study did not generate new unique reagents.

### Data and code availability

- Raw, preprocessed data, and experimental program have been deposited at an view-only OSF repository (https://tinyurl.com/4y844ymh) and will be made public upon publication.
- All original analysis code and study stimuli has been deposited at Github and is publicly available at https://github.com/YueyingDong/yd2026_paper as of the date of publication.

## ACKNOWLEDGMENTS

We thank Tim Brady for his input on stimulus processing and task design. We also thank members of the Kiyonaga lab for helpful comments on the manuscript: Vivien Chopurian, Pria Daniel, Lana Gaspariani, Janna Wennberg, Connie Xie, Zhuojun Ying, and Sihan Yang. This work was supported by NIH award 1R01EY036843-01 to A.K.

## AUTHOR CONTRIBUTIONS

Conceptualization, A.K. and Y.D.; methodology, Y.D. and Y.H.; Investigation, C.X., Y.H., and Y.D.; writing—original draft, A.K. and Y.D.; writing—review & editing, A.K. and Y.D.; funding acquisition, A.K.; resources, A.K. and Y.D.; supervision, A.K.

## DECLARATION OF INTERESTS

None.

## DECLARATION OF GENERATIVE AI AND AI-ASSISTED TECHNOLOGIES

During the preparation of this work, the author(s) used Claude AI in order to refactor and provide documentation for the analysis code. After using this tool or service, the author(s) reviewed and edited the content as needed and take(s) full responsibility for the content of the publication.

## STAR★METHODS

### METHOD DETAILS

#### Participants

The experiment was approved by the University of California, San Diego’s Office of IRB Administration (OIA). We recruited 46 adult participants (over 18 years of age), with normal or corrected to normal (via contact lens) vision, through UCSD’s SONA platform. Related studies that tested a brightness-related pupillary effect in WM have included 20 - 30 participants^2,27^. In our previous study (36 participants, 7791 trials) and its replication (49 participants, 13864 trials), we reliably observed this WM pupillary effect^16^. Therefore, here we aimed for a similar sample size.

Participants completed informed consent and received course credit as compensation. Trials were removed either for poor eyetracking data quality (see Preprocessing for Eyetracking QA details) or a missing behavioral response. Participants were excluded if they had more than 40% of trials rejected, across the QA criteria. After excluding 2 participants, 44 remained in the final analyses (*34 female, ages 18-26*). The study took 60 - 80 minutes to complete.

#### Task procedures

The experiment was programmed and presented in Psychopy, v2023.2.3. Participants completed the task in a dimly lit testing room, on a 1920 x 1080 monitor (24 inch, 60.96cm) with a 60hz refresh rate. Participants sat 60cm away from the screen center, their heads stabilized by a chinrest. We implemented continuous eye tracking throughout the experiment via a desk-mounted Eyelink Portable Duo (SR Research Ltd.), recording at 1000hz. We performed a 13-point calibration using the built-in calibration module, and performed a drift check at the beginning of every block (every 20 trials) to ensure that central fixation was maintained.

The WM stimuli were photographs taken during either daytime or nighttime, drawn from 9 categories: airport, bench, circus tent, ferris wheel, gas station, house, playground, school bus, and shopping cart. Images were processed to remove identifying information (e.g., words or people), grayscaled, and luminance-matched within each the day and nighttime conditions (see *Image Processing* for sample, probe, and manipulation check image specifications). Images were then cropped to 256 * 256 pixels, subtending 6.68º visual angle on the screen.

Each trial started with a fixation period of 1000ms, during which an empty circle appeared at screen center. The circle’s RGB color value ([128, 115, 96]) was equiluminant to the neutral grey background (RGB value: [128,128, 128]). Next, two WM sample images appeared on either side of the circle for 3000ms, each centered at ±5.64º visual angle. The sample array always contained one daytime and one nighttime scene from different categories, and the location of the day-vs. night-time scene was counterbalanced across trials. The images were followed by noise masks for 250ms and then a 250ms fixation period. Then a retrocue appeared for 500ms, comprising a half-filled semicircle signaling which image would be tested at the end of the trial (e.g., if the filled portion were on the right, the image from the right of center would be tested). Then followed a 6000ms delay after which the probe array appeared for 5000ms. Participants saw six probe images and the task was to click on the one that matched the cued WM sample.

Critically, we manipulated which feature of the cued stimuli (visual detail or semantic category) was relevant for the memory probe. The task-relevant feature was indicated by the color of the retrocue (either green or purple). On “visual” trials, participants had to select the probe image that was an exact match to the cued image. On “semantic” trials, participants had to select the image that was from the same category as the cued image. The mapping of green (RGB [0,128,0]) and purple (RGB [128,0,128]) retrocue colors to the visual and semantic test conditions was counterbalanced across participants. The type of images in the probe array also differed accordingly for the visual and semantic trials. For both conditions, probe arrays were constructed to include images from 3 base categories – the correct cued category, the uncued category, and an unrelated category – as well as corresponding lures for each of those 3 categories.

On visual trials, the probe array included the two WM sample stimuli, an image from one of the remaining sample stimulus categories, and an additional lure selected from each of these three categories (See *Stimulus image processing* for all categories). For example, if the WM sample images were a circus tent and a playground (as shown in **Figure 1**), then the probe would include these two images, one different circus tent, one different playground, and two unrelated images (e.g. two school buses). If the circus tent had been flagged as task-relevant by the retrocue, the correct probe response would entail clicking the specific cued circus tent.

On semantic trials, the probe array included a new image from the same semantic category as each of the two WM samples, another image from one of the remaining sample stimuli categories, and lures to correspond to each of these categories. For example, if the WM sample images were a circus tent and a children’s playground, then the probe would include images of a different circus tent and a different children’s playground, as well as an image from a third category (e.g. a school bus). Then lures from visually similar categories were selected for each of those categories. For example, the lures for a circus tent, children’s playground, and school bus would be a camping tent, a bouncy house, and an ambulance, respectively (See **Table 1** for lure category mappings). On semantic trials, neither of the exact WM sample images reappeared in the probe display. If the circus tent had been retrocued, the correct response would entail clicking on the new circus tent image.

**Table 1.** WM sample stimulus categories (used in all conditions) and lure categories (used in the probe array on semantic trials).

| Sample categories | Lure categories |
| --- | --- |
| House | Barn |
| Bench | Porch swing |
| Playground | Bounce house |
| School bus | Ambulance |
| Circus tent | Camping tent |
| Shopping cart | Wheelbarrow |
| Airplane | Helicopter |
| Gas station | Toll booth |
| Ferris wheel | Swing ride |

Therefore, the probe lures on *visual* trials were meant to be semantically similar to the WM content, so that a strong visual representation would be required for successful performance. The lures on *semantic* trials, on the other hand, were meant to be visually similar to the WM content, so that a clear semantic representation would be necessary for successful performance (see *Image Processing*). The screen locations for all probe images were randomly intermixed. Note that all probe images were highpass filtered to remove the luminance feature, rendering the daytime vs. nighttime element task-irrelevant for identifying the correct item.

In theory the retrocue and probe manipulation should encourage participants to use a more visual WM maintenance strategy when visual detail is required, and a more semantic maintenance strategy when a category label is required. To ensure that participants engaged in the task as expected, we also incorporated a manipulation check on 30% of the trials. On these trials an intervening task appeared in the middle of the delay period, involving either a visual or a semantic judgement (counterbalanced across *visual* and *semantic* WM trials). In the visual manipulation check, participants were shown an image and needed to judge whether it was scrambled or intact. For the semantic manipulation check, participants saw a word string and needed to judge whether it described a man-made or natural object. In both cases, they used a keyboard button press to indicate their response (‘w’ for scrambled/manmade, ‘d’ for intact/natural). Manipulation check tasks appeared 3000ms into the WM delay, remained on the screen for 1500ms, and then were followed by a blank 1500ms delay before the probe. Thus the total trial length was equated on all trials. If participants were using the retrocues as designed, then visual manipulation checks should be relatively more disruptive to WM during *visual* cue trials, while semantic manipulation checks should be more disruptive during *semantic* cue trials.

The experiment comprised 216 trials in total, broken into 18 blocks of 20 trials each. 108 stimulus images were drawn from 9 image categories, with 12 images per category; 6 daytime scenes and 6 nighttime scenes for each category. Each unique sample stimulus appeared twice in the experiment, once on the left and once on the right side of fixation. All participants completed at least 20 trials of practice before the experiment. After the experiment, participants completed a 16-item Vividness of Visual Imagery Questionnaire (VVIQ^34,76^), as a self-report index of their visual imagery.

#### Stimulus image processing

All WM sample images were photographs taken during day and nighttime, collected from Wikipedia Commons and keyword searches, according to the categories in Table 1. We manually removed all identifiable information, such as faces or identifying text, using Photoshop (Adobe Inc.). All images were then grayscale and cropped to 256 * 256 pixels. We created 39-52 unique images in each sample category for the total image set, and then selected an equal number from each category to include in the task.

We created two copies of the images, one to be processed as WM sample stimuli, and the other for their inclusion in the probe array (**Figure 5A**). For the sample stimulus set, we applied luminance matching within the daytime and nighttime scenes separately, using the MATLAB (The Mathworks, Inc.) **SHINE** toolbox to equate the brightness histograms^77^. The mean luminance for the day- and nighttime scenes was set to 70% and 25% on the HSV scale, respectively. The probe images did not require luminance matching, because they were high-pass filtered to remove luminance information. In addition to the sample images, we also collected “lure” images for inclusion in the probe arrays. These images were drawn from related categories to the sample categories (see **Table 1** for stimulus and lure categories), and were also grayscaled and high-pass filtered to remove luminance.

**Figure 5:**
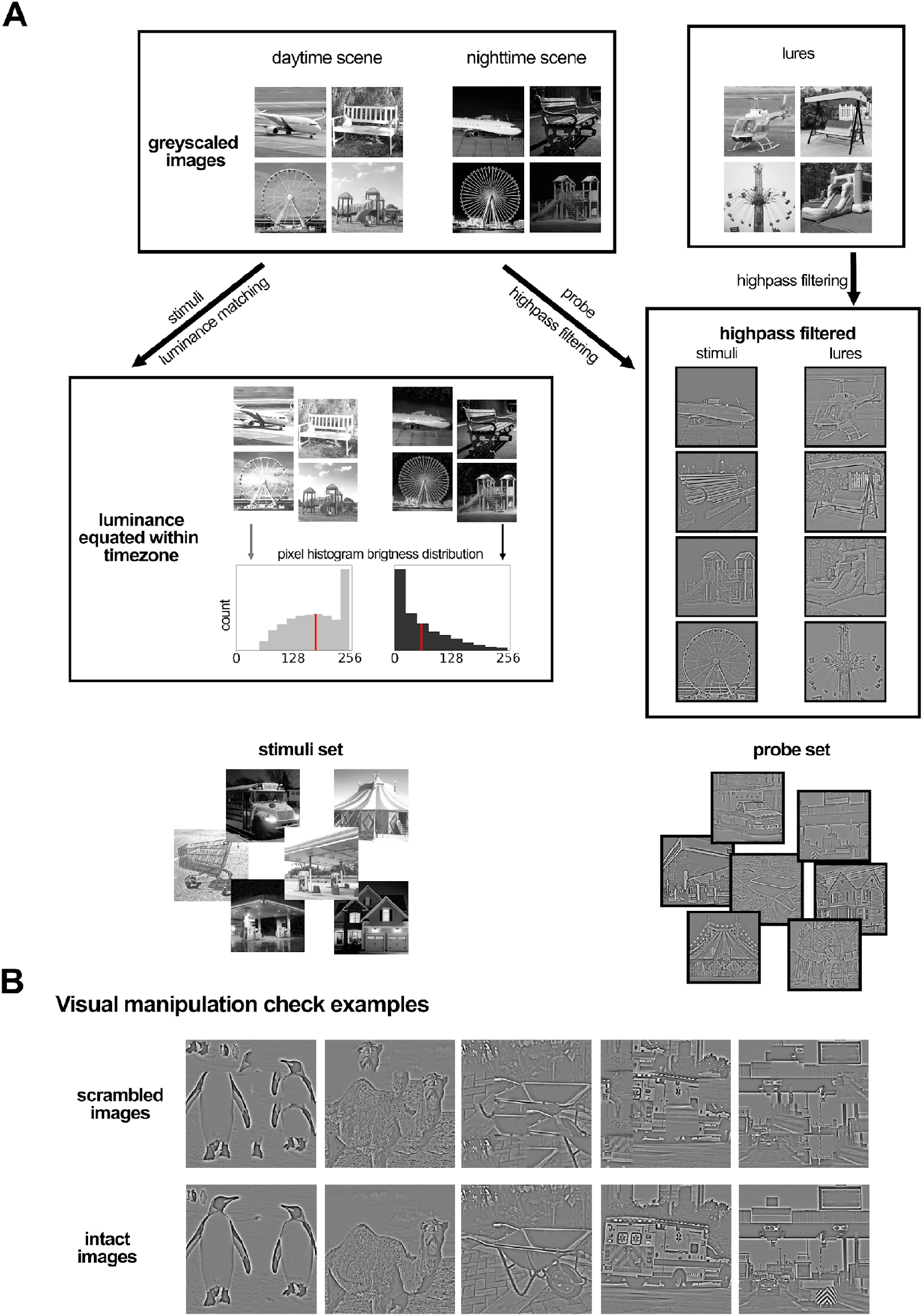
Sample stimulus and probe processing procedure. (**A**) The WM samples and probes were curated from the gray-scaled image set. The WM samples were luminance-equated within each day and night groupings; the probes were high-pass filtered to remove luminance. (**B**) Example scrambled vs. intact stimuli used for visual manipulation checks.

We aimed for the “visual lures” to be semantically similar to the WM samples but visually distinct, and for the “semantic lures” to be visually similar to the WM samples but semantically distinct. We therefore implemented additional probe selection criteria to match the lures across visual and semantic trials. We quantified visual similarity between probe pairs using the Structural Similarity Index (SSIM), a commonly used metric to compare images in terms of brightness, contrast, and structure^78^. Then for each WM sample stimulus, the *visual* lures were randomly drawn from the bottom 30%, least similar-looking images within the category. On the other hand, the *semantic* lures were drawn from the lure category, filtered for the top 30%, most similar-looking images.

We also created additional stimulus sets for the manipulation check task. For the visual manipulation check, we collected a set of images consisting of half manmade and half natural objects. In addition to the semantic lure images, we gathered images from nine natural object categories (Bamboo; Camel; Dandelion; Bird’s nest; Platypus; Shark; Starfish; Tulip; Penguin), drawn from THINGS database^79^. After greyscaling and high-pass filtering, we scrambled the images by cutting them into 16 tiles, randomly swapped 30% of the tiles, and applied a gaussian blur to the edges of each flipped tile (**Figure 5B**). We therefore created two versions of each image scrambled or intact), from 18 unique categories, totaling 36 stimuli.

For the semantic manipulation check, the words were drawn from one of the 36 categories listed in **Table 2**. There were 18 categorical labels for the visual manipulation check, with an intact and a scrambled image for each category, totaling 36 stimuli. Therefore, we added another 18 words to the semantic manipulation check stimulus set, 9 natural objects and 9 manmade objects. We simplified some category terms (e.g. replacing ‘camping tent’ with ‘tent’, ‘swing ride’ to ‘carousel’) to remove semantic ambiguity.

**Table 2.**
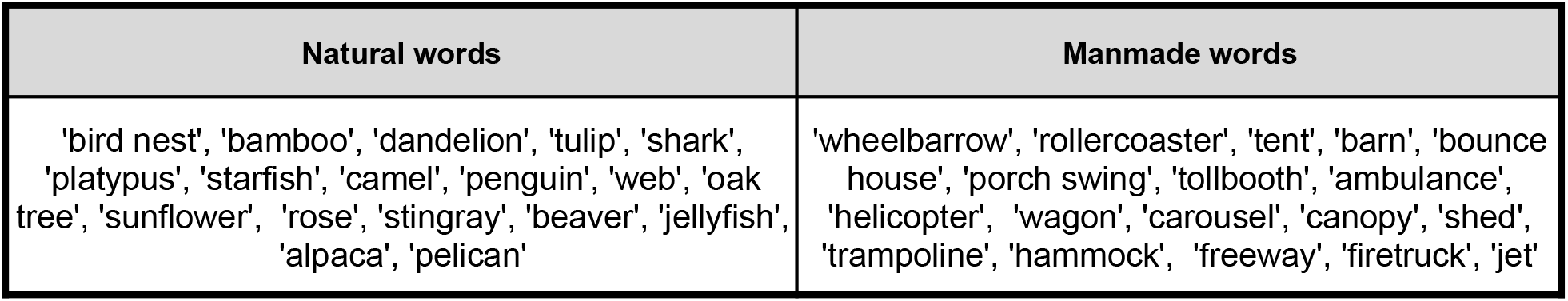
Semantic manipulation check terms from natural and manmade objects.

| Natural words | Manmade words |
| --- | --- |
| 'bird nest', 'bamboo', 'dandelion', 'tulip', 'shark', 'platypus', 'starfish', 'camel', 'penguin', 'web', 'oak tree', 'sunflower', 'rose', 'stingray', 'beaver', 'jellyfish', 'alpaca', 'pelican' | 'wheelbarrow', 'rollercoaster', 'tent', 'barn', 'bounce house', 'porch swing', 'tollbooth', 'ambulance', 'helicopter', 'wagon', 'carousel', 'canopy', 'shed', 'trampoline', 'hammock', 'freeway', 'firetruck', 'jet' |

#### Eyetracking data processing

##### Preprocessing

The eye-tracking processing pipeline closely followed procedures established in previous work^16,43^. Data was exported using the EyeLink Data Viewer software package (Version 4.3.1).We detected blink artifacts as outliers in the velocity space of pupillary change. The data were first transformed into *Trial x Time* format, and for each trial, we obtained a smoothed derivative array, 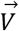, from the original pupil size, 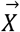

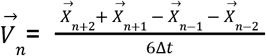

Next, we defined blinks as any time point that exceeded a velocity threshold

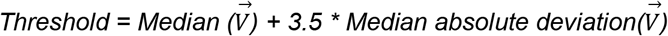

The blinks were first identified as any period that exceeded this threshold, and then padded with a 50ms buffer on each side. This blink mask was applied to both pupil and gaze data, rejecting any timepoints during these blinks. For each gap, depending on the availability of data surrounding each gap, we used cubic spline or linear interpolation to reconstruct the pupil, and nearest-neighbor or linear for gaze. If a gap exceeded 2 seconds, then no interpolation was attempted.

After interpolation, a trial was discarded if it had more than 10% data missing from the main epoch of interest – that is, the 3500 ms from retrocue onset to memory delay offset. After participant exclusion, trial-wise data cleaning retained 93.3% of data in the experiment (149 - 216 trials per participant)

##### Pupil analysis

For every trial, we calculated an individual’s baseline as the average pupil size over the 200ms prior to retrocue onset. We subtracted this baseline value from our epoch of interest, which included the retrocue period (500ms) and the first half of the delay (3000ms). We excluded the second half of the delay because the manipulation check appeared on 30% of the trials during this epoch.

As in previous work^16,43^, we used permutation *t-*tests to identify periods of significant differences between the conditions of interest. To do this, we downsampled the data to 10 Hz by averaging the data points in every 100-ms time bin. This served to smooth the time series and render the permutation testing more computationally efficient. On each permutation iteration, we randomly shuffled the brightness labels (daytime/nighttime). Then for each time point, we applied an independent *t-*test between the pupil size in the shuffled day-versus nighttime categories. We identified clusters of significance and summed up the *t* values corresponding to the largest cluster, yielding a *t*_*sum*_ in each permutation repeat. This process was repeated 5000 times to generate a null distribution. Lastly, we conducted the same independent *t-*test on the real data with unshuffled labels to identify the clusters of significance. We compared the *t*_*sum*_ of each cluster to the null distribution, and only registered those that exceeded the 95th percentile as significant. We ran this procedure separately for the visual and semantic trials.

##### Saccade analysis

The processing procedure and parameters were all derived from prior studies^43,80^. The blink mask was identified from the pupil data and subsequently used to filter blink artifacts on the gaze data. Following artifact rejection, the missing data were interpolated with either linear or nearest neighbor interpolation, depending on the availability of nearby data points.

Next, we extracted saccades from this preprocessed gaze data. Similar to how blinks were identified as outliers in the speed of pupil change, saccades were characterized as outliers in the speed of gaze dislocation^40^. We defined a minimum dislocation threshold of 8 pixels (0.21° visual angle) to prevent false positive characterizations. Additionally, if a trial was deleted from the pupil data following the procedures in *Preprocessing*, it was also rejected from the gaze data.

We characterized every identified saccade by its direction relative to the cue. If the horizontal dislocation of a saccade was in alignment with the cued hemifield, then this saccade was marked as “toward”. Otherwise, a saccade would be marked as “away”. For example, if the left side was cued, we would only characterize a leftward change in the horizontal coordinate of gaze as “toward”.

##### IPA analysis

The Index of Pupillary Activity (IPA) is a measure of pupil diameter oscillations that is meant to be an indicator of cognitive load^36^. The algorithm we used was adapted from Duchowski et al., 2018. It performs a Discrete Wavelet Transformation (DWT) on the pupil signal and identifies the number of prominent frequencies.

Unlike the main pupil analyses, after performing artifact rejection, we didn’t interpolate the missing data. For every trial, we first measured the overall IPA using the pupil data during the retrocue and delay epochs. Then, to gauge the time course of effort deployment, we applied the IPA algorithm over a 100ms rolling window over the data.

## Notes

### Competing Interest Statement

The authors have declared no competing interest.

### Summary of Updates

Updated/revised introduction and discussion in accordance with the reviewer's suggestions.

